# Single cell RNA-sequencing identifies the gene expression profile of the asexual and sexual invasive merozoites of the human malaria parasite

**DOI:** 10.1101/2024.09.26.614664

**Authors:** Laura Piel, Celine Hernandez, Yan Jaszczyszyn, Delphine Naquin, Lucille Mellottée, Joana M. Santos

## Abstract

The intraerythrocytic developmental cycle (IDC) during which the malaria parasite *Plasmodium falciparum* multiplies asexually initiates when parasite stages called merozoites are released into the bloodstream, following schizont rupture, and invade the host red blood cells. The merozoite is the less studied stage of the IDC, despite its importance for the establishment of infection. Here we uncovered the gene expression profile of the free merozoites, by doing single cell RNA-sequencing (scRNA-seq). We found that free merozoites express invasion and motility genes that are actively transcribed in schizonts and after merozoite egress. *P*. *falciparum* parasites undergo at least one IDC before sexually-committing and transforming into the sexual stages, the gametocytes, which are then transmitted to the mosquito vector. Our scRNA-seq dataset identified two early states of sexual commitment, preceding expression of the gametocytogenesis master regulator *ap2-g*. We hypothesize that early committed parasites are primed for sexual differentiation and definitive engagement into gametocytogenesis occurs when *ap2-g* is expressed. Our results contribute to a better understanding of how the parasite prepares for invasion and how it exits asexual growth and enters sexual differentiation. These transitions ensure survival of the parasite in the human host and transmission to the mosquito vector so the gene markers identified here can be therapeutically explored in the future to block malaria invasion or transmission.

**Importance:** The malaria parasite *Plasmodium falciparum*, which causes the death of over half a million people each year, has a complex life cycle but symptoms only arise when the parasite invades and develops inside the host red blood cells. The merozoites are the parasite stages responsible for invading the human red blood cells and they are one of the few extracellular stages of the lifecycle, even if briefly. However, their gene expression profile had yet to be determined. Here we used single cell RNA-sequencing to establish the gene expression program of free merozoites. We also identified for the first time the gene signature of sexually-committed and sexual merozoites that will later give raise to gametocytes and ensure parasite transmission.

## Introduction

Malaria, responsible for over half a million deaths each year (1), is an infectious disease caused by the unicellular eukaryotic parasite *Plasmodium*. Out of the five species of parasite that cause disease in humans, *P*. *falciparum*, is the most virulent. The parasite life cycle is complex, alternating between the human host and the mosquito, that acts as both a host and a vector. The intraerythrocytic developmental cycle (IDC), in which the parasite multiplies within the human host red blood cell (RBC) for 48-hours, starts when parasite stages called merozoites, invade the host RBC (Figure 1A). Once inside the RBC, the parasite transforms into rings, then trophozoites and finally schizonts. At the end of schizogony, new merozoites are released into the bloodstream and the IDC restarts. In *P. falciparum*, the IDC ensures parasite multiplication but also transmission because parasites must undergo at least one IDC in order to sexually commit and form gametocytes that are later transmitted to the mosquito vector (2) (Figure 1A). Sexual commitment only occurs in a small subset of cells (<10%).

**Figure 1.**
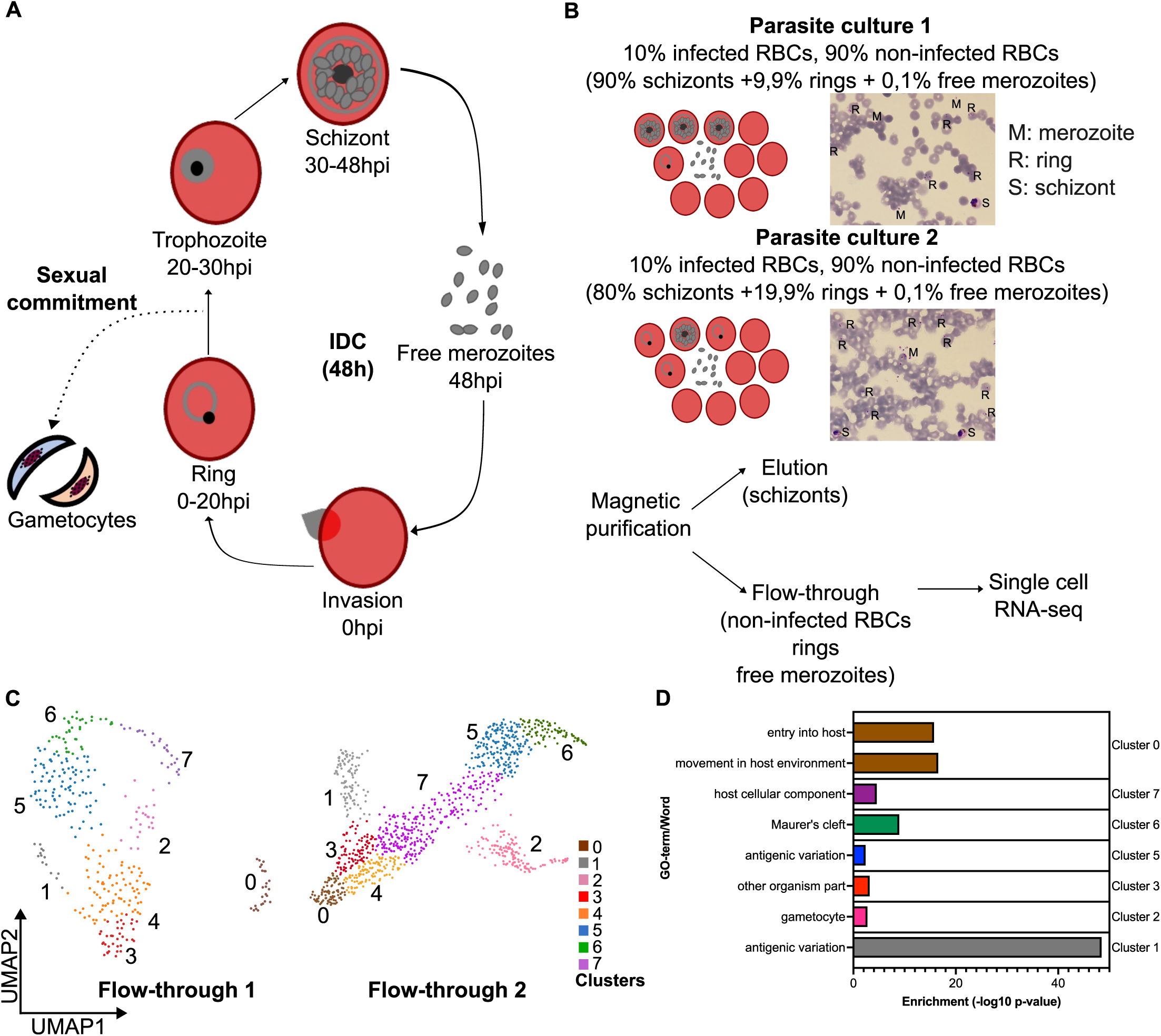
Single cell RNA-sequencing of merozoites. (**A**) During the intraerythrocytic developmental cycle (IDC), lasting 48 hours, parasite forms called merozoites egress from schizonts and invade the red blood cell (RBC) to become first rings and later on trophozoites. These transform into schizonts, within which new merozoites are formed. Parasites can also sexually commit, leaving the IDC, eventually differentiating into gametocytes. The merozoites are only briefly extracellular, after schizont egress and before RBC invasion. Numbers below each stage indicate hours post-invasion, hpi. (**B**) Parasites were magnetically purified from two different cultures containing 10% infected RBCs. As determined by Giemsa-stained blood smears, the first parasite culture contained 90% schizont-stage parasites, 9.9% ring-stage parasites and 0.1% free merozoites and the second one 80% schizont-stage parasites, 19.9% ring-stage parasites and 0.1% free merozoites (R: rings, S: schizonts, M: merozoites). The schizonts were retained in the magnetic columns and in the flow-through, we collected the non-infected RBCs, the ring-stage parasites and the free merozoites. The flow-through fractions from the two parasite cultures were processed independently for single cell RNA-seq using 10x Genomics (the number of sequenced cells is in Table S1). (**C**) Uniform Manifold Approximation and Projection (UMAP) of the flow-through cells, from the two parasite cultures, colored according to the Seurat cluster. The number of cells on each cluster is in Table S1 and the code used for the Seurat analysis is in the supplementary files 1 and 2. (**D**) GO-term analysis of the marker genes of each flow-through cluster. Because there is no GO-term associated with gametocytes, word search analysis was done instead for cluster 2 genes. Only some GO-terms/words are shown. The list of marker genes as well as of all GO-terms and words is in Table S2.

The merozoite is the less studied stage of the IDC, despite its attractiveness as a drug target given that: (i) it is extracellular, even if briefly; (ii) it is responsible for RBC invasion, which is indispensable for parasite survival; and, (iii) it initiates the IDC, which is the symptomatic life cycle stage of malaria. Studies describing the gene expression profiling of the different stages of the *P. falciparum* life cycle, lack information for the merozoite stage (3) and, it is only recently that it was shown that merozoites are transcriptionally active (4). RBC invasion is a fast process, lasting ∼1 minute (5), and merozoites only last 5-15 minutes extracellular (6), so extracellular merozoites have never been isolated and characterized.

Here we used single cell RNA-sequencing to establish for the first time the gene expression signature of individual free merozoites, upon schizont rupture. We find that free merozoites express invasion and motility genes that are actively transcribed in schizonts and merozoites. We also identify two early states of sexual commitment that prepare the parasite to enter gametocytogenesis and be transmitted.

Our comprehensive study further sheds light into gene expression in the malaria parasite and unfolds for the first time the single cell expression atlas of the human malaria invasive merozoites, allowing the parasite to invade the RBC.

## Results

### Identification of free merozoites

*Plasmodium falciparum* merozoites have so far been isolated for transcriptional studies by forced rupture of purified schizonts, after parasite treatment with an egress inhibitor that inhibits merozoite egress (4). The recovered merozoites are viable but short-lived (6). In order to record the gene expression profile of *de facto* free merozoites, we decided to not block egress and this way allow natural merozoite release from schizonts. To separate the free merozoites from the schizonts, we used magnetic columns that retain the trophozoites and schizonts (6), which are paramagnetic due to the accumulation of hemozoin, the end product of hemoglobin digestion (Figure 1B). The free merozoites, as well as the rings and non-infected red blood cells, are not retained by the magnetic column and are then recovered in the flow-through of the magnetic purification (Figure 1B). Merozoites cannot be separated from the other cells of the flow-through so, to identify the gene expression profile of the free merozoites, we used single cell RNA-sequencing (scRNA-seq), which is powerful in identifying discrete cell types from mixed cultures.

To enrich for the presence of free merozoites in the parasite culture, we single synchronized parasites for two cycles, and when schizonts were visible in Giemsa-stained smears, we collected the culture. We used two different parasite cultures. One containing mainly schizonts (parasite culture 1) and one containing a mix of schizonts and rings (parasite culture 2) (Figure 1B). This strategy was fruitful because free merozoites were identified in both cultures in Giemsa-stained blood smears (Figure 1B). We passed each parasite culture, independently, through magnetic columns, and the purity of the elution and flow-through fractions was then verified by blood smear (Figures S1A-B). The two flow-through fractions were then prepared individually for scRNA-seq using the 10X Genomics technology.

After sequencing, we obtained reads from 1464 flow-through cells, 405 from the first parasite culture, and 1059 from the second parasite culture (Table S1). We acquired 200 million reads, with a high percentage of sequencing saturation (over 98%) (Table S1). On average, each cell expressed on mean 600 genes (Table S1, Figures S1C-D). After filtering out cells with fewer than 200 genes and over 2500 genes per cell, a common threshold to analyze *P*. *falciparum* single cell data (3,7,8), we retained 1421 cells in the combined flow-through fractions so 97% of the initial sequenced cells (Table S1).

In agreement with what we had observed in blood smears (Figure 1B), the majority of flow-through cells are ring-stage parasites because they express the ring marker *ring-exported protein 2* (*rex2*, PF3D7_0936000) (Figure S1C). 2-4% of cells, however, express the merozoite marker *merozoite surface protein 1* (*msp1*, PF3D7_0930300) (Figures S1E-G) and are putative merozoites. We used *msp1* as a merozoite marker because MSP1 is the most abundant merozoite surface protein (9) and it is over-expressed in merozoite cells versus ring cells, as determined by Reers *et al*. (4), that defined a list of genes that are over-expressed in merozoites versus rings or schizonts.

Merozoites and ring-staged cells are expected to express different genes so we used Seurat, a toolbox for quality check, analysis and visualization of single cell data (10–12), to non-biasedly identify a cell cluster, in the flow-through fraction, corresponding to free merozoites (Supplementary files 1 and 2). Seurat identified eight expression clusters (Figures 1C and S1H-I, Table S2). Differential expression analysis was done to determine which genes are over-expressed in one particular cluster versus the others. These genes are herein called marker genes. Gene ontology (GO) analysis of the marker genes of each flow-through cluster shows that cluster 0 cells express genes specialized in host cell entry and motility (Figure 1D, Table S2), functions associated with invasion, and so with merozoites. Furthermore, 95% of these genes were previously shown to be over-expressed in merozoites (4) and 44% of them encode proteins previously identified in merozoite proteomes (13,14) (Figure S1I, Table S2). Additionally, as expected for merozoites, the cluster 0 cells contain less RNA than other flow-through cluster cells (Figure S1J). This indicates that the cluster 0 cells are free merozoites. We identified 113 free merozoite cells in the two flow-throughs (Table S1), corresponding to less than 10% of the population, as anticipated from an elusive stage that had never before been captured by RNA-sequencing studies.

### Free merozoites express genes encoding proteins implicated in invasion and motility

We identified 102 genes over-expressed in free merozoites versus the other cells of the flow-through (Figures 2A-B, Table S2). The most differentially expressed genes are those encoding the erythrocyte binding antigen 175 (EBA-175), the gamete release protein (GAMER), the reticulocyte binding protein homologue 1 (Rh1), the calcium dependent protein kinase 5 (CDPK5) and the myosin A-tail interacting protein (MTIP) proteins (Figure 2A). CDPK5, Rh1 and EBA-175 have all been shown to play a role in discharge of the parasite apical organelles, micronemes and rhoptries. These Apicomplexa-specific organelles store proteins that are released into the parasite surface prior to invasion (15). CDPK5 and Rh1 are critical for micronemes discharge (16,17). The micronemes store, amongst others, EBA-175. EBA-175 is a parasite ligand that establishes a specific interaction with the host cell receptor glycophorin-A (18). Establishment of this interaction triggers rhoptry discharge (19). Release of the rhoptry proteins requires proper positioning of the organelle, which is under the responsibility of the armadillo-domain containing rhoptry protein (ARO) (20), and *aro* is a marker gene of free merozoites (Figures 2A-B, Table S2).

**Figure 2.**
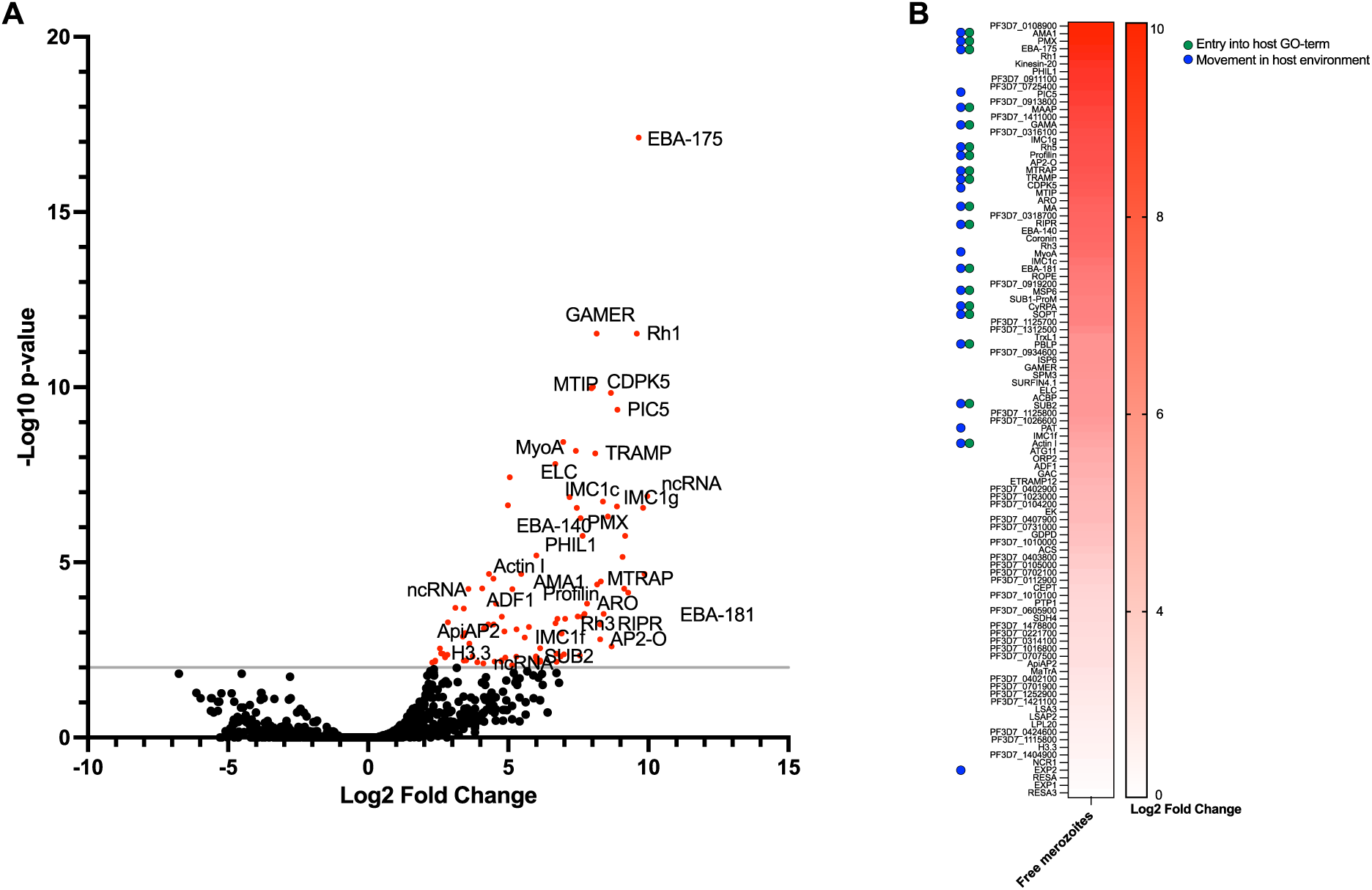
Gene signature of the free merozoites. (**A**) Volcano plot of the genes expressed in free merozoites. Genes overexpressed in these cells, with a p-value equal or less than 0,01, versus other cells of the flow-through, are in red. Some of the gene names are shown. The complete list of genes is in Table S2. (**B**) Heatmap showing the log2 fold change expression value of the marker genes of the free merozoites. If the genes code proteins implicated in entry into the host or in motility, according to GO-term analysis, is indicated by a green or blue dot, respectively. The log2 fold change for each gene and the full GO-term analysis is in Table S2 and Figure S2A.

After strong attachment of the parasite to the RBC surface, via the establishment of the ligand-receptor interactions, the parasite reorientates (21). This requires photosensitized INA-labeled protein (PHIL1) (22) and its partner protein PIC5 (23), and both coding genes are over-expressed in free merozoites (Figures 2A-B, Table S2). Afterwards, the parasite ligand Rh5 interacts with the host basigin, and there is formation of the tight junction. Free merozoites over-express the genes coding for Rh5 and its interacting partners RIPR and Cysteine-Rich Protective Antigen (CyRPA) (24), as well as apical membrane antigen 1 (AMA1) (Figures 2A-B, Table S2), member of the tight junction (21). Rh5 only binds basagin, when proteolytically processed by Plasmepsin X (PMX) (25), and *pmx* is also a signature gene of free merozoites (Figures 2A-B, Table S2).

Malaria merozoites glide prior to invasion (26). This is supported by the parasite actomyosin machinery, the glideosome (Figure S2B), which also propels the parasite inside the host (27). Free merozoites over-express several of the genes encoding glideosome constituents (27): Myosin A (MyoA), MyoA-tail Interacting Protein (MTIP), Myosin Essential Light Chain (ELC) and Glideosome-Associated Connector (GAC) (Figures 2A-B and S2B, Table S2). We also detect expression of the genes coding MTRAP and TRAMP, which might interact with GAC (26), and several genes coding for inner membrane complex 1 (IMC1) proteins, responsible for anchoring and stabilizing the glideosome (27) (Figures 2A-B and S2B, Table S2). Gliding requires actin polymerization by profilin (27) and we detect over-expression of the *profilin*, *actin I* and *actin depolymerizing factor 1* genes in extracellular merozoites (Figures 2A-B and S2B, Table S2).

The gene signature of free merozoites reflects then their invasive and motile nature (Figures 2B and S2A, Table S2). Intriguingly, these cells also over-express three non-coding RNAs (Table S2). Among them, *pf3d7_0108900*, an antisense RNA covering the 5’ untranslated region of the *phil1* gene (28), which, as mentioned, is a marker gene of free merozoites. There are also seventeen genes coding for proteins of unknown function that may also be implicated in invasion or motility (Table S2).

Tripathi *et al.* showed variation in gene expression between merozoites in schizonts of the same age and established a metric called variability index score (VIS) to reflect this variation (8). We determined the VIS of each of the free merozoites signature genes, by comparing our data to theirs. Most free merozoite genes have a VIS of 1 or more (Figure S2C, Table S2). That is not the case of the marker genes of other clusters (Figure S2C), suggesting that the free merozoite genes in particular are variably expressed among schizonts.

### Free merozoites genes are transcribed during schizogony and after merozoite egress

Merozoites have been shown to be transcriptionally active (4) so we wondered if the free merozoites marker genes are actively transcribed in merozoites. To access this, we first determined the time of peak transcription of each marker gene, as established by a published real-time expression data (29). 54% of genes are peak transcribed at 1 or 48 hpi, when the merozoites prepare to invade/egress and so when they are extracellular or are about to (Figure 3A, S3 Table). 37% of genes reach peak transcription beforehand, during schizogony, between 30 and 47hpi, and the remaining 9% are transcribed following invasion, in rings (2-20hpi) or trophozoites (20-30hpi) (Figure 3A, Table S2). In *P*. *falciparum*, active gene promoters are labelled by the histone marks H3K9ac, H3K27ac and H3K4me3 (30), so we examined if these marks are present in the promoters of the free merozoites signature genes. According to previous Chromatin ImmunoPrecipitation (ChIP-Seq) data (4), the three histone marks are present in the gene promoters, as well as in the transcription start sites, in merozoites and schizonts (Figure 3B), but not in ring-staged parasites (Figure S3A). Binding then drops within gene bodies (Figure 3B). These two comparative analyses indicate that the free merozoite genes are transcribed during schizogony and when the merozoites are extracellular. Bulk RNA-seq (4) indicated that 89% of the free merozoites marker genes are more expressed in merozoites than schizonts so peak expression of the marker genes may be reached after merozoite egress. In ring-staged parasites, H3K27ac is removed from all genes, except from those to be expressed in rings (31), so these analyses also show that the free merozoite genes are no longer transcribed after invasion. This is further supported by the fact that only one of the free merozoites marker genes is, according to (4), over-expressed in rings versus merozoite parasites (Figure S3B, Table S2).

**Figure 3.**
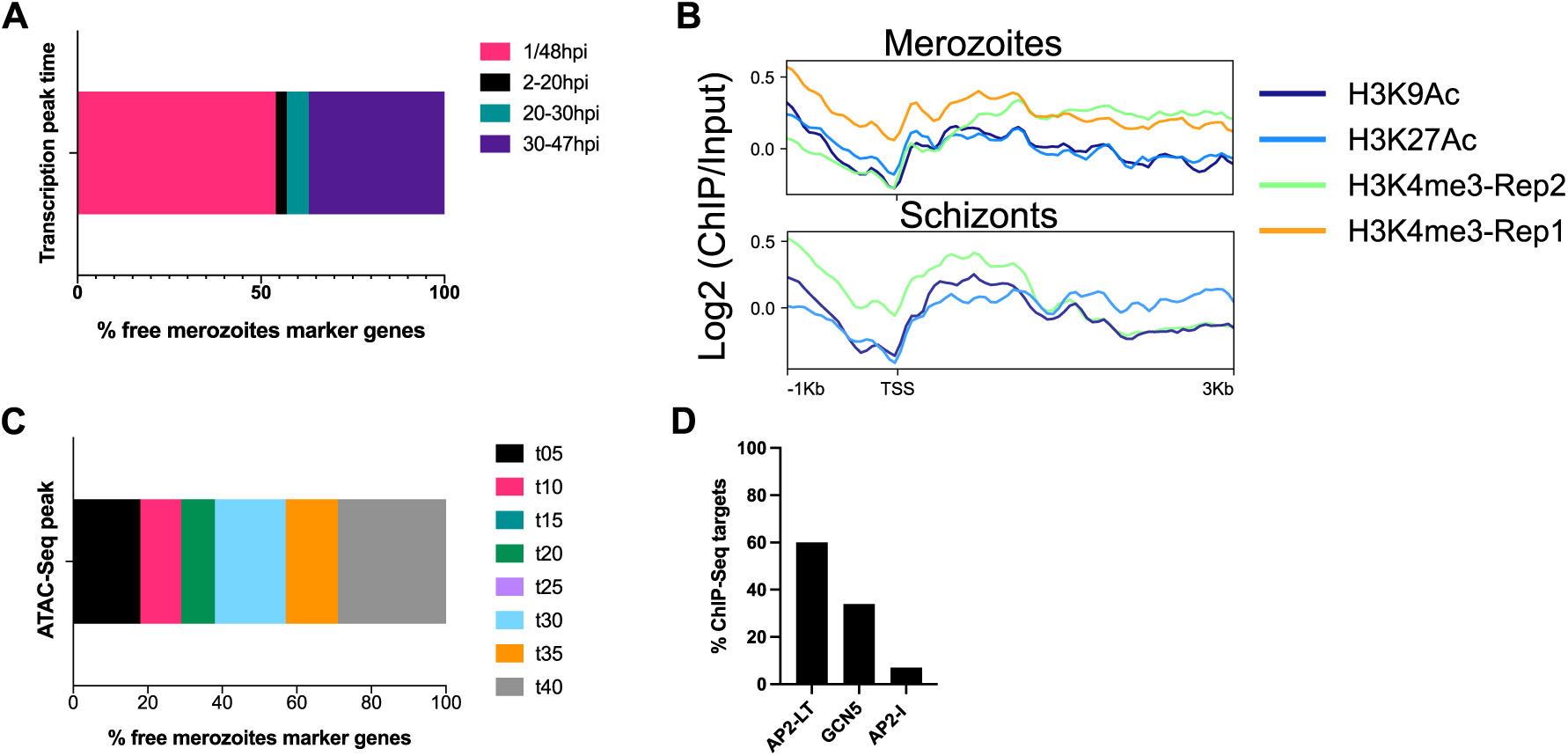
Free merozoite genes are transcribed in schizonts and merozoites. (**A**) As determined by Painter *et al.* (29), most marker genes of free merozoites are peak transcribed in schizonts (30-47hpi, purple) or merozoites (1/48hpi, pink), whereas a minority are peak transcribed in rings (2-20hpi, black) or trophozoites (20-30hpi, green). The timing of peak transcription for each gene is in Table S2. (**B**) Graph-plots showing binding of the histone marks H3K9ac (dark blue), H3K27ac (light blue) and H3K4me3 (green and orange) to the gene promoters, transcription start sites (TSS) and gene bodies of the free merozoite genes, in schizont and merozoite parasites, as determined by the log2 ChIP/Input calculated by Reers *et al*. (4). Binding in rings is in Figure S3A. (**C**) According to the ATAC-Seq dataset of Toenhaken *et al.* (32), most free merozoites gene promoters are euchromatic in schizonts (t30, t35 and t40 in blue, orange and gray, respectively), while less than 40% are euchromatic in rings (t05, t10 and t15 in black, pink and sage, respectively) or trophozoites (t20 and t25, in in green and purple, respectively). The ATAC-Seq peak for each gene is in Table S2. (**D**) Bar graph showing how many free merozoite genes are ChIP-Seq targets of AP2-LT, GCN5 or AP2-I, according to previous datasets (35–39). Which genes are ChIP-Seq targets of these proteins is in Table S2.

H3K4me3 is a marker of open, accessible, chromatin (30), so we examined if the promoters of the free merozoites marker genes are accessible when the genes are being transcribed, i.e., in schizonts and merozoites. We did this by comparing our dataset to a published time-course of chromatin accessibility in *P*. *falciparum* (32). In that study, 62% of the free merozoites gene promoters are euchromatic between 30 and 40hpi, so during schizogony (Figure 3C, Table S2). This analysis suggests that chromatin accessibility precedes gene transcription in free merozoites. This points towards activation of gene transcription by a transcription factor.

The ApiAP2 proteins are the main family of DNA-binding proteins in the Apicomplexa and several have been shown to act as transcription factors (33). We examined expression of *apiap2* genes in free merozoites to identify proteins potentially implicated in regulating expression of the free merozoite genes. We detected expression of 16 *apiap2* genes, including *ap2-lt* (Figure S3C). AP2-LT has been previously found to bind the promoters of several invasion genes (34,35) and to interact with the histone acetylase GCN5, which binds to H3K9ac-labeled promoters (36,37). Since we showed that the free merozoites express invasion genes and these genes promoters are marked by H3K9ac, we determined how many of the free merozoites promoters are bound by AP2-LT or GCN5, according to published ChIP-Seq datasets (34–37). GCN5 binds 40% of the promoters and 60% of these are bound by AP2-LT, suggesting that AP2-LT and GCN5 may regulate expression of the free merozoites genes (Figure 3D, Table S2). On the contrary, AP2-I, which has also been shown to regulate expression of some invasion genes (38) and it is expressed by free merozoites (Figure S3C), only binds upstream of 7% of the free merozoites genes according to (38,39) (Figure 3D, Table S2).

### Identification of an early sexual commitment state by unsupervised clustering

To increase the power of our scRNA-seq dataset, we integrated the two flow-through datasets and applied Seurat to it (Figures 4A and S4A, Supplementary file 3). This led to the identification of 13 clusters (Table S3). Then we used the list of signature genes of free merozoites, that we previously established (Table S2), to find free merozoite cells in the integrated dataset. Cells expressing the free merozoite marker genes are found in clusters 4 and 13 (Figures 4B and S4B). These cells contain less RNA and encode less transcripts than other cells (S4C Figure), express the merozoite marker *msp1* (Figure S4D) and encode invasion-related genes (Figure 4C, Table S3), which are all hallmarks of extracellular merozoites. On the other hand, the other cells of the integrated dataset express the *rex2* ring marker gene and are therefore rings (Figure S4E).

**Figure 4.**
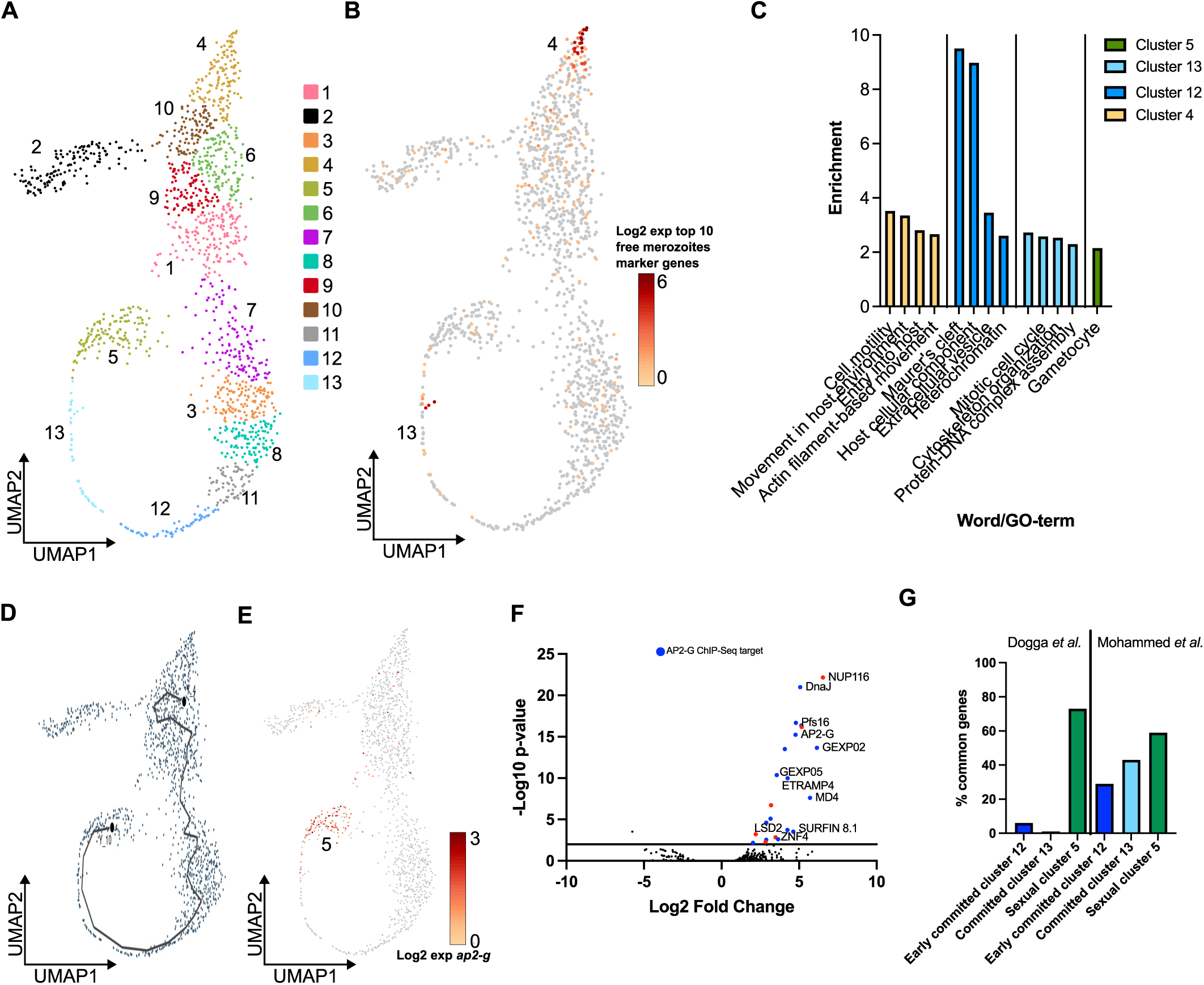
Identification of early sexually-committed and sexually-committed cells. (**A**) Seurat clusters of the integrated flow-through dataset (the full list of genes expressed in each cluster is in Table S3, the number of cells on each cluster is in Table S1 and the Seurat analysis are in supplementary file 3). (**B**) There are two sub-groups of free merozoites, one in cluster 4 and one in cluster 13, as determined by the expression of the top 10 free merozoite markers genes. (**C**) GO-term analysis of the marker genes of clusters 4, 5, 12 and 13 showing that cluster 4 cells express invasion-related genes and are thus asexual, cluster 12 cells code exported proteins, cluster 13 cells express DNA replication, chromatin and cell-cycle genes, while cluster 5 cells express gametocyte-related genes (the full GO-term analysis are in Table S3). Bars are colored according to the cluster. (**D**) Pseudotemporal analysis of the merozoite clusters showing that there is only one trajectory. The cells pseudotime is in Table S1 and the Monocle analysis is in supplementary file 3. (**E**) UMAP representation showing that cells in cluster 5 express *ap2-g*, indicating that they are destined to become gametocytes and are therefore sexual. (**F**) Volcano plot of the genes expressed in cluster 5 cells versus the other cells of the integrated dataset. Genes overexpressed with a p-value equal or less than 0,01 are in red. AP2-G ChIP-Seq targets, as determined by Josling *et al*. (39), are in blue. The names of selected genes are shown. The full list of genes is in Table S3. (**G**) Graph showing that many of the signature genes of cluster 5 cells (green) were previously reported by Dogga *et al*. (7) or Mohammed *et al*. (42) as being up-regulated in committed cells or early gametocytes, respectively. Several of the marker genes of cluster 12 (dark blue) or cluster 13 cells (light blue) were also detected in the Mohammed *et al*. dataset (42) (the list of common genes between datasets is in Table S3). Bars are colored according to the Seurat cluster.

The few marker genes in common between the clusters 4 and 13 cells are signature genes of free merozoites (Figure S4F), indicating that the cells in the two clusters correspond to two distinct sub-populations of merozoites. To allocate an identity to these sub-populations, we did trajectory analysis. Cluster 4 cells have the earliest pseudotime (Figures 4D and S4G, Table S1) and express invasion and motility genes (Figure 4C, Table S3) so we conclude that these 141 cells are asexual parasites. 17% of these cells are free merozoites (Table S1). We propose that the other cells are very early rings that still express the gene expression program of free merozoites.

The cells trajectory terminates in cluster 5 (Figures 4D and S4G). Cluster 5 parasites express gametocyte-related genes (Figure 4C, Table S3), namely *ap2-g* (Figure 4E), the master regulator of gametocytogenesis (40,41), indicating that they have entered sexual commitment. This is further supported by the fact that, of the 35 marker genes of cluster 5 parasites (Figure 4F, Table S3), 25 were previously identified as markers of sexually-committed cells (7,42) (Figure 4G, Table S3) and 57% are ChIP-Seq targets of AP2-G (39) (Figure 4F, Table S3). The cells of clusters 12 and 13, preceding cluster 5, do not express *ap2-g* (Figure 4E). They do, however, over-express genes previously shown to be expressed in gametocyte precursor cells by another scRNA-seq study (42). 29% of the cluster 12 marker genes and 43% of the cluster 13 marker genes (Figure 4H, Table S3) were shown by that study to be expressed by sexually-committed parasites, before bifurcation of the cells into the male and female gametocyte trajectories (42). This indicates that clusters 12 and 13 precede sexual differentiation. We used the definition of Dogga *et al*. (7) that defined the beginning of sexual commitment as the moment when parasites enter the sexual trajectory and here name the cluster 12 parasites, early sexually committed, and the cluster 13 parasites, sexually committed. To distinguish these parasites from the ones in cluster 5, we herein call the cluster 5 parasites, sexual. We have then identified two early states of sexual commitment, preceding expression of the *ap2-g* master regulator. The early commitment state (cluster 12) is defined by the expression of 34 marker genes, the commitment state (cluster 13) by the expression of 242 marker genes, and there are 22 signature genes of sexual parasites (cluster 5) (Table S3). We identified 60 early sexually-committed parasites, 46 committed parasites and 130 sexual parasites (Table S1), corresponding altogether to 17% of the flow-through population, as expected since few parasites exit asexual growth and sexually-commit.

### Early sexually committed and committed cells have distinct gene expression signatures

We compared the gene expression profile of the early committed (cluster 12) and committed parasites (cluster 13) with that of the asexual and the sexual parasites (cluster 4 and cluster 5, respectively) to identify differences among them. Apart from the free merozoites marker genes, that as noted before, are also expressed in committed parasites (Figure S4F), marker genes of asexual parasites are under-expressed in early committed and committed parasites (Figure 5A, Table S3). Similarly, marker genes of early committed and committed parasites are down-expressed in asexual parasites (Figures 5B-C, Table S3). Further supporting that asexual and sexually-committed parasites have distinct gene expression profiles, several of the sexual marker genes of cluster 5 are already expressed in sexually-committed parasites, but not in asexual cells (Figure 5D, Table S3). That is the case of *gametocyte exported protein 2* (*gexp02*), *male development protein 4* (*md4*), *parasitophorous vacuole membrane protein 16* (*pfs16*), *surface-associated interspersed protein 8.1* (*surfin8.1*) and *early-transcribed membrane protein 4* (*etramp4)* (Figure 5D, Table S3). This concords with previous studies that showed that some early sexual markers are transcribed in the absence of AP2-G (43–46).

**Figure 5.**
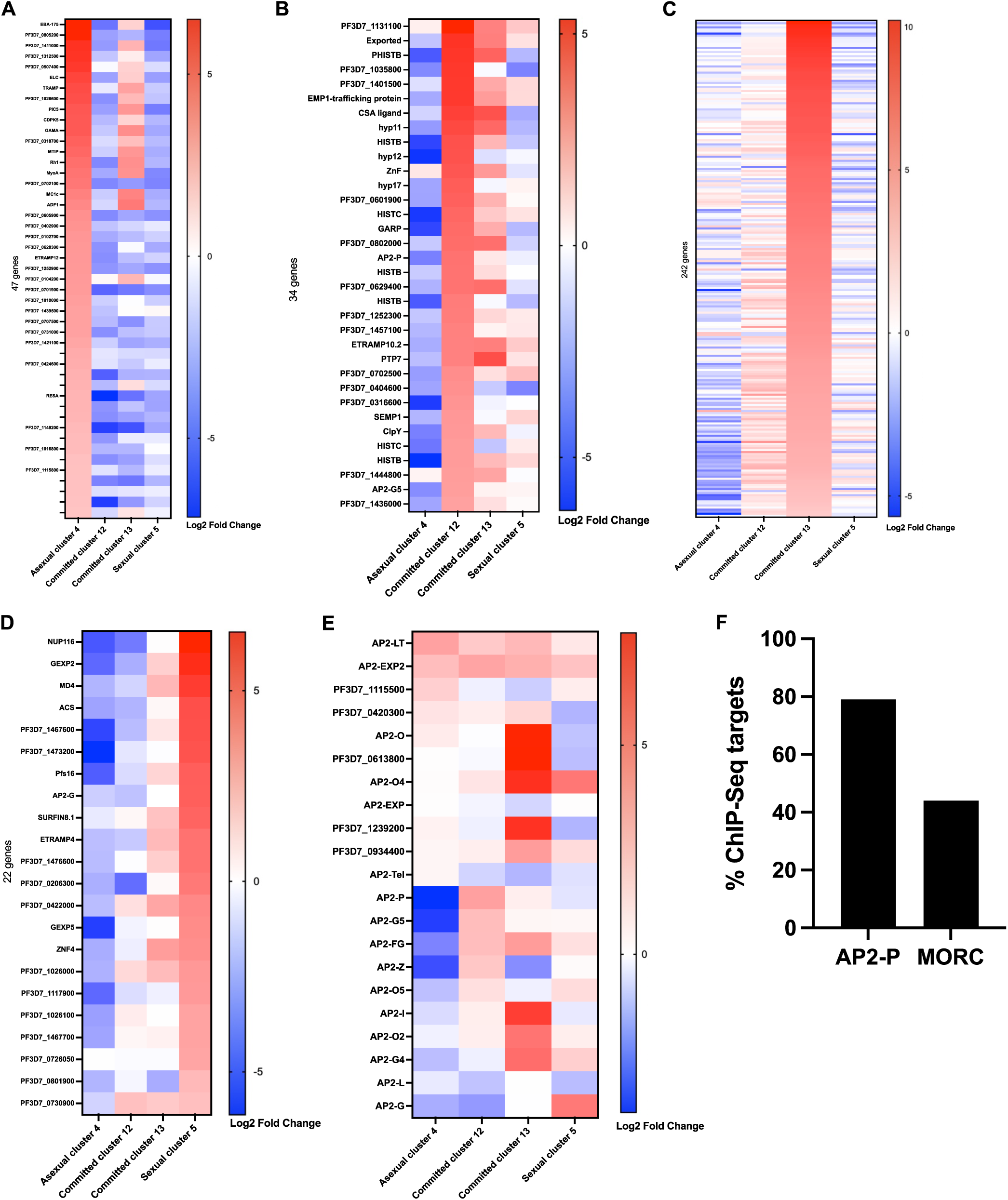
Comparison of the early sexually-committed and sexually-committed cells. (**A**) Heatmap showing the log2 expression levels of the signature genes of asexual cells (cluster 4) on those cells as well as in the early committed, committed and sexual cells (cluster 12, cluster 13 and cluster 5, respectively), showing that, apart from the free merozoite gene markers, which are also expressed in committed cells, asexual cells have a distinct gene expression profile from other cells. Annotated rows correspond to genes expressed in free merozoites. (**B**) Heatmap showing the log2 expression levels of the signature genes of early committed cells (cluster 12) on those cells as well as in the asexual, committed and sexual cells (cluster 4, cluster 13 and cluster 5, respectively), showing that some of those genes are expressed in committed and sexual cells but not in asexual cells. (**C**) Heatmap showing the log2 expression levels of the signature genes of committed cells (cluster 13) on those cells as well as in the asexual, early committed and sexual cells (cluster 4, cluster 12 and cluster 5, respectively), showing that several of those genes are already expressed in early committed cells. (**D**) Heatmap showing the log2 expression levels of the signature genes of sexual cells (cluster 5) on those cells as well as in the asexual, early committed and committed cells (cluster 4, cluster 12 and cluster 13, respectively), showing that many of these genes are already expressed in committed cells. The log2 expression data for **A**, **B**, **C** and **D** is on Table S3. (**E**) Heatmap showing the log2 expression levels of *apiap2* genes in asexual, early committed, committed and sexual cells (cluster 4, cluster 12, cluster 13 and cluster 5, respectively). (**F**) Bar graph showing how many early committed genes are ChIP-Seq targets of AP2-P or MORC, according to previous datasets (50–53). Which genes are targets is in Table S3.

Many of the marker genes of early-committed and committed parasites encode ApiAP2 DNA-binding proteins (Table S3) and committed parasites also over-express genes coding for zinc finger and RNA-binding proteins (Figure S5B, Table S3). This is compatible with the initiation of a distinct gene expression program that prepares the parasite to exit asexual growth and transform into the sexual stages.

Early committed genes are still expressed in sexually-committed parasites, and vice-versa, but each cell type has a distinct gene expression profile (Figures 5B-C). GO-term analysis of the marker genes of clusters 12 and 13 show that early committed cells encode proteins to be exported to the host cell cytosol and membrane, and early committed cells code for cytoskeleton and DNA replication proteins (Figure 4C, Table S3).

We detect transcripts for female and male markers in early committed and committed parasites, suggesting, like previous studies (42), that some sex-specific transcripts are expressed early on during sexual commitment. Namely, *ap2-g5*, linked to the male lineage (42), is over-expressed by early committed parasites, and *ap2-fg*, *ap2-o* and *ap2-o2*, associated with the female lineage (42,47), are marker genes of committed parasites (Figures S4A-B, Table S3).

To identify potential regulators of sexual commitment, we determined which *apiap2* genes are expressed in asexual (cluster 4), early committed (cluster 12), committed (cluster 13) and sexual (cluster 5) parasites (Figure 5E). *ap2-lt* is highly expressed in asexual cells and it continues to be expressed during sexual commitment (Figure 5E). As mentioned, AP2-LT may regulate expression of the free merozoites genes but its expression in early gametocytes was also reported (42), so it is likely that, like other ApiAP2 proteins, AP2-LT regulates multiple steps of the parasite lifecycle. *pf3d7_0420300* has the same expression pattern as *ap2-lt* (Figure 5E) and its expression during gametocytogenesis was also recently put forward (42). Expression of *ap2-exp* and *pf3d7_1115500* is higher in asexual and sexual parasites than during sexual commitment (Figure 5E). AP2-EXP regulates expression of genes encoding exported proteins (35,48) so this might reflect a specific need of the asexual and sexual parasites to export proteins to the host cell cytosol and surface. AP2-G5 represses gametocytogenesis by suppressing expression of *ap2-g* (49) and its transcription decreases in sexual cells (Figure 5E).

*ap2-p*, which has been linked to female gametocyte development (42), is a marker gene of early committed parasites (Figures 5B and S5A, Table S3), so we used two published ChIP-Seq datasets (50,51), to determine how many of the early commitment gene promoters are bound by AP2-P. Strikingly, 79% of the cluster 12 marker genes are AP2-P ChIP-Seq targets (Figure 5F, Table S3). AP2-P is implicated in chromatin boundary insulation, together with MORC (51), and, according to previous ChIP-Seq assays (51–53), some of the early committed gene promoters are also bound by MORC (Figure 5F, Table S3). Our results then suggest a role for AP2-P in regulating early sexual commitment, perhaps in complex with MORC. *ap2-z*, which is transcribed in gametocytes but only translated in the mosquito, is also highly expressed by early committed parasites (Figure 5E).

Six *apiap2* genes are markers of committed cells: *ap2-i*, *pf3d7_1239200*, *pf3d7_0613800*, *ap2-fg*, *ap2-o* and *ap2-o2* (Figure S5B, Table S3). Three others, *ap2-o4*, *ap2-g4* and *pf3d7_0934400*, are highly expressed in these cells (Figure 5E). *ap2-o*, *ap2-i*, *ap2-g4* and *pf3d7_1239200* have previously been shown to be expressed in early gametocytes (42) so they may play a role in the early steps of sexual commitment. This is further supported by the fact that AP2-I interacts with AP2-G (39) and that AP2-G4 plays a positive regulatory role in gametocytogenesis (49). However, comparison of published ChIP-Seq datasets of AP2-I and AP2-G4 (38,39,49), with the list of marker genes of committed cells, only identified a few target genes (data not shown).

*ap2-g* is only expressed in sexual parasites (Figure 5E) and it is a marker gene of these cells (Figures 4F and 5D, Table S3). Since *ap2-g* is transcribed mainly at the ring and late schizont stages (45), it is not surprising that we detect its expression in our dataset of rings and merozoite parasites. Sexual parasites also express *ap2-fg* and *ap2-o2* (Figure 5E) that are expressed in mature gametocytes (42,47).

### Early sexually-committed rings can be produced by next cycle conversion

We identified one sexual free merozoite and three sexually-committed merozoites but we were unable to identify early sexually-committed free merozoites (Table S1). Because *P*. *falciparum* parasites can sexually-commit at the ring-stage by same cycle conversion (SCC), without activation of sexual commitment, prior to invasion (45), we wondered if the ring-stage parasites we labelled as early committed are produced by SCC. This would explain why there are no early committed merozoites. Alternatively, the early sexually-committed rings were generated, upon RBC invasion of early committed merozoites (next cycle conversion, NCC), and we did not identify early committed merozoites due to reduced cell numbers. There are minor transcriptional differences between sexually-committed parasites produced by NCC or SCC (45) so to explore this, we searched for intracellular parasites expressing the early commitment genes. To do that, we passed the schizonts that had been retained in the magnetic columns through a filter (6) and prepared them for single cell RNA-sequencing. We obtained a similar number of reads, sequencing saturation and reads mapped to the genome in the intracellular cells, as compared to the flow-through cells (Table S1). After filtering with the same parameters used for the flow-through cells, we retained 191 intracellular cells (Table S1).

To discover intracellular cells expressing early-committed genes, we integrated the data for the 191 intracellular cells with the combined flow-throughs, and applied Seurat to this new integrated dataset (Supplementary file 4). We identified again 13 clusters (Figures 6A and S6A, Table S1). Cluster 11 cells express early commitment genes, parasites expressing commitment genes are in cluster 10, and cluster 7 cells, succeeding cluster 10, express sexual marker genes (Figure S6B, Table S4). 8 of the 66 early committed parasites and 24 of the 69 committed parasites, are intracellular (Figure 6B, Table S1). Since we identified intracellular parasites expressing early commitment and commitment genes, we conclude that early committed ring-staged parasites can be generated by NCC. This does not exclude that they can also be generated by SCC.

**Figure 6.**
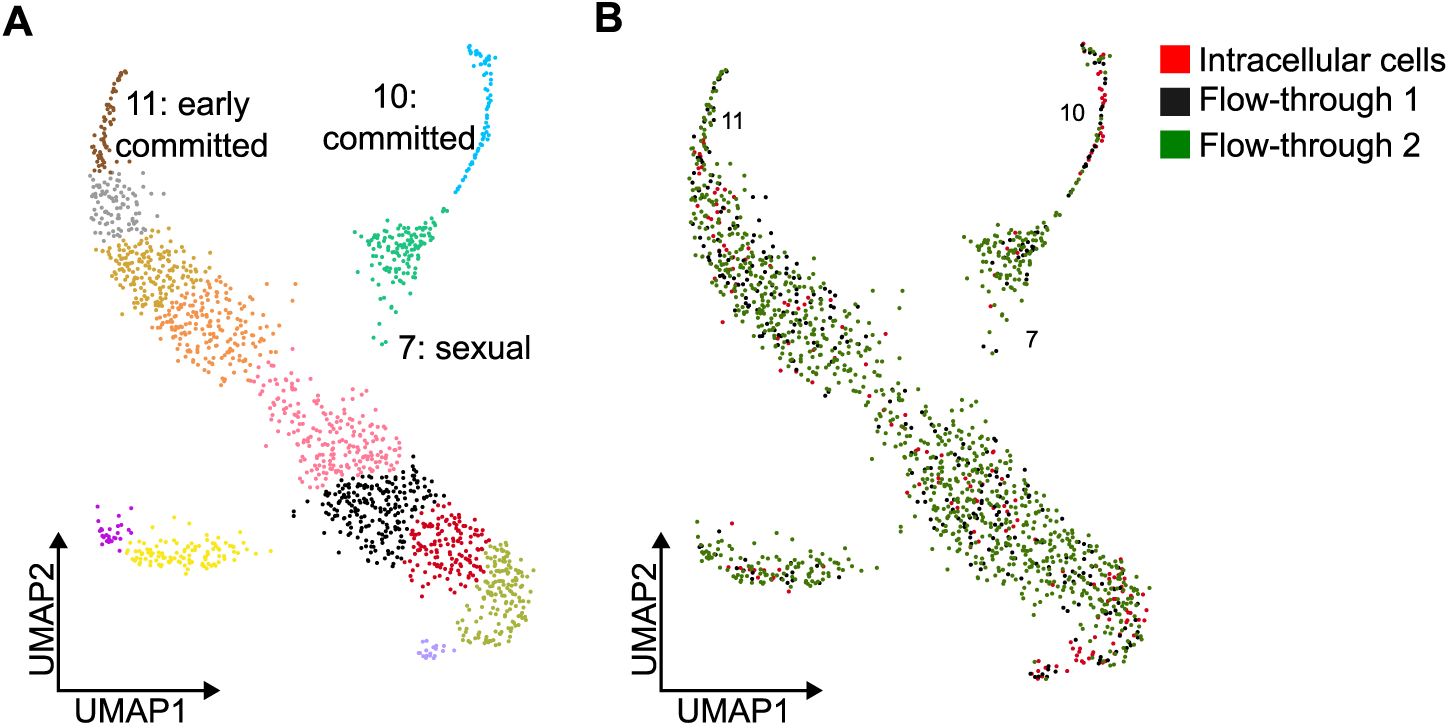
Early sexually-committed and sexually-committed genes are expressed by intracellular parasites. (**A**) UMAP plot showing which Seurat clusters of the integrated dataset of the flow-through and intracellular cells correspond to early committed, committed and sexual cells. The number of cells in each cluster is in Table S1 and the Seurat analysis are in supplementary file 4. The list of genes expressed by each cluster is in Table S4. The other cluster numbers are in Figure S6A. (**B**) UMAP plots showing the number of intracellular cells (red) or flow-through cells (black and green) on each cluster. The number and the type of cells (intracellular, flow-through) in each cluster is in Table S1.

## Discussion

Here we used single cell RNA-sequencing to identify the gene expression profile of the RBC invasive stages of the malaria parasite, the merozoites. We validated these cells identity by showing that they express the merozoite marker *msp1*, that several of the free merozoite genes identified here encode previously identified merozoite proteins (13,14) and that almost all of the marker genes of these cells were previously shown to be over-expressed in merozoites (4).

Extracellular merozoites express invasion and motility genes, as expected from cells that need to glide and invade the host. The promoters of the free merozoites signature genes are already marked with active histone marks during schizogony and continue to be so at merozoites (4), suggesting that the genes are actively transcribed in the two developmental stages. This is further supported by the genes peak transcription time, according to a previous study (29). After invasion, the genes are no longer transcribed. Bulk RNA-seq showed that invasion genes are transcribed during schizogony (54–56) but here we show, by doing comparative analysis, that these genes are also transcribed in free merozoites.

Sexual merozoites had never been identified but since *P*. *falciparum parasites* must invade RBCs before sexually committing (2), these had to exist. We identified these rare cells as well as sexually-committed parasites. We propose that parasites enter sexual differentiation when they express *ap2-g* and thus called the cells expressing *ap2-g*, sexual. These sexual cells express many of the genes expressed by what Dogga *et al*. called committed cells (7). Here we show that two gene expression commitment programs, that we call early sexual-commitment and sexual-commitment, precede expression of the sexual genes. The clusters corresponding to the early sexually-committed and committed parasites were defined by Seurat, using standard parameters, and were thus found in an unsupervised manner. Each cluster contains several cells and it is characterized by the expression of many marker genes. There are 242 marker genes of sexually-committed cells and 34 markers of early committed cells. In comparison, there are 22 marker genes of sexual cells. Intracellular parasites also express early sexual commitment and commitment genes so expression of these genes is not restricted to free merozoites and rings. Early sexually-committed and committed trophozoites must also exist but we did not have this parasite stage in our cultures. Even though many of the genes expressed by what we here call early committed and committed cells were previously identified as being expressed in early gametocytes (42), the existence of an early sexual commitment state was not reported. We think that in our work we were able to detect more slight changes in gene expression and this way distinguish between early sexually-committed, committed and sexual cells, because we only analyzed two parasite stages, while other studies used mixed stages.

We propose that the early sexual commitment steps identified here correspond to an early stage of sexual commitment that primes parasites for sexual differentiation. Definitive commitment into gametocytogenesis would then require *ap2-g* expression. Further studies are required to test this hypothesis.

*P*. *falciparum* parasites can sexually differentiate at the ring-stage, without expressing AP2-G in the previous IDC via the SCC pathway (45). This occurs when AP2-G protein levels reach a threshold sufficient in rings to trigger the sexual differentiation program (45). It is unknown how many of the ring parasites expressing *ap2-g*, of our dataset, also express the AP2-G protein, but since only 7 of the 134 sexual cells are intracellular and, as mentioned, only one of those cells is a free merozoite, it is possible that most of the sexual rings we identified were produced by SCC, as proposed by Bancells *et al.* (45).

We propose that AP2-LT regulates expression of the free merozoites genes and that AP2-P may regulate the early commitment expression program because many of the marker genes of free merozoites or early commitment parasites are ChIP-Seq targets of AP2-LT or AP2-P (34,35,50,51), respectively. AP2-LT is part of the SAGA activating complex, together with GCN5 (36,37,57,58), and GCN5 also binds upstream of several free merozoite genes (36,37). *ap2-lt* is also expressed during sexual differentiation, as shown here and in a previous study (42), so this protein may have multiple roles. The precise role of AP2-LT in regulating gene expression requires the establishment of a conditional knockout or knockdown, since gene disruption seems impossible (33).

AP2-P, which may or may not regulate expression of the early committed genes, establishes chromatin boundaries, together with MORC, and specific ApiAP2 proteins (51). While many early committed gene promoters are bound by AP2-P (50,51), fewer are bound by MORC (51–53), so it is unclear if AP2-P associates with MORC during sexual commitment. AP2-P has been previously characterized but its implication in sexual commitment was not explored (51–53). The existing AP2-P conditional knockout was done in a parasite strain that is a poor gametocyte-producer (50), so to study the function of the protein in early sexual commitment, a new transgenic line in a more gametocyte prone strain, would have to be established. Alternatively, one could use the glmS conditional knockdown system but this system does not induce complete protein shutdown (51).

In summary, we identified here the gene expression profile of extracellular merozoites, as well of the early stages of sexual commitment, despite the rarity of these cells. This study reveals how the parasite prepares for invasion and how it exits asexual growth and enters sexual differentiation. These transitions ensure survival of the parasite in the human host and transmission to the mosquito vector so the gene markers identified here can be therapeutically explored in the future to block malaria invasion or transmission.

## Materials and Methods

### Parasite culturing and merozoites purification

*P. falciparum* W2mef parasites were cultured at 37°C in the presence of 5% oxygen and 5% carbon dioxide, in RPMI 1640 media (Fisher) supplemented with hypoxanthine and 0.5% Albumax II (Invitrogen) in 2% hematocrit. Blood was obtained from the Etablissement Français du Sang (EFS).

Parasite synchronization to the ring stage was done by incubating pelleted cultures 1:1 with 0.3M L-Alanine, 10mM HEPES pH 7.5 solution for 10 min at 37°C, after which cultures were centrifuged, resuspended in fresh media and placed back in culture (59).

For scRNA-seq, two mixed parasite cultures of non-infected RBCs, schizonts, rings and free merozoites were passed through two LD columns (Miltenyi Biotec), washed with RPMI culture and eluted with RPMI into Falcon tubes. The flow-through, corresponding to the ring stage parasites, free merozoites and non-infected red blood cells, was also recovered. The elution corresponds to the schizont fraction. Schizonts were passed through a 1.2µM filter (Pall) to release the intracellular cells inside (6). After centrifugation, the intracellular or the flow-through cells were counted with a Neubauer chamber and diluted with 1XPBS/0.04%BSA to 1 million cells/ml before loading into the Chromium.

The purity of each fraction was verified by Giemsa-stained blood smear. Images of blood smears were acquired with the widefield microscope Eclipse (Nikon) at 60x amplification.

To stain with DAPI, purified schizonts were fixed in 1XPBS/4%formaldehyde, washed and deposited into glass slides mounted with VectaShield Antifade medium containing DAPI (Vector laboratories) and imaged with a SP8 confocal microscope (Leica) at 63x amplification. Images were treated with ImageJ.

### Single cell RNA-Sequencing

Approximately 10.000 cells (intracellular or flow-through) were loaded on a Chromium Chip G (10x Genomics) as per the manufacturer’s recommendations. Chips containing cell suspensions, gel beads in emulsion (GEMs) and reverse transcription reagents were loaded into the chromium controller (10x Genomics) for single cell partitioning and cell barcoding. Barcoded cDNA and sequencing libraries were generated according to the Chromium Next GEM Single Cell 3’ gene expression v3.1 protocol. Quantification and the quality of sequencing libraries was verified on an Agilent Bioanalyzer. Pooled libraries were sequenced on a NextSeq 2000 system (Illumina).

### Data analysis

Quality control checks on the raw sequencing data were performed using FASTQC (60). FastQ files were analyzed using the CellRanger software (10X Genomics), including alignment, filtering and quantitation of reads on the *Plasmodium falciparum* genome (3D7) and generation of feature-barcode count matrices. During mapping, the reads corresponding to the human host cell were discarded.

Genes detected in fewer than three cells were removed. Cells with more than 0.5% of mitochondrial genes were removed. Cells with fewer than 200 genes and more than 2500 genes per cells were removed.

The Seurat toolbox version 4.3.0.1 was used to normalize each dataset using SCTransform (61) and cluster the cells in an unsupervised manner, using standard conditions (all code used is in the supplementary files). To determine if the cells recovered in the flow-through of the second magnetic purification express the same expression clusters as the ones recovered in the flow-through of the first purification, we looked for cells expressing marker genes of each of the flow-through clusters and then manually identified them as belonging to that cluster. Visualizations and identification of differentially expressed genes were done using the Loupe browser (10X Genomics, version 7).

Graphs showing the binding pattern of the active histone marks H3K9ac, H3K4me3 and H3K27ac in the gene promoters and gene bodies of the free merozoites were generated using computeMatrix and plotProfile from the deepTools package (62) of Galaxy (63) using the ChIP-Seq data from.

Pseudotime and trajectory analysis were calculated using Monocle3 R package version 1.3.4 (64) (thecode used is in the supplementary file 3).

The ATAC-Seq analyses were done by comparing the genes expressed in each cluster to those identified in (32). Whenever more than one peak was assigned to a gene promoter, we only considered the earlier time point. For most genes, there was only one peak of accessibility.

The transcription peak time of the marker genes expressed in each cluster was calculated in (29).

To determine the percentage of genes encoding proteins detected in mature merozoites by mass spectrometry we compared our dataset to the high-throughput LC-MS/MS of *P*. *falciparum* 3D7 merozoites data generated by Shahid Kahn from Leiden University that identified over 1000 proteins (13) or to the data of Kumar *et al*. (14). We only considered proteins detected with at least 2 peptides.

The ChIP-Seq comparison analyses were done by determining if the genes expressed in a specific cluster were identified as target genes in (34–39,48,50–53).

The percentage of genes over-expressed in merozoites versus rings or schizonts was determined by comparing our dataset to that of (4).

Venn diagrams were generated using the tool BioVenn (65).

Gene Ontology (GO) enrichment analysis was done using the GO feature at PlasmoDB (release 68) (66).

Graph plots were generated with GraphPad Prism version 10.0.0 for Mac, GraphPad Software, Boston, Massachusetts USA, www.graphpad.com.

## Data availability

All RNA sequencing data generated in this study has been deposited in NCBI’s Gene Expression Omnibus and is accessible through GEO series accession number GSE279639.

## Code availabity

All code used in this study is presented in the Supplementary files.

## Acknowledgements

This work was funded by a CNRS ATIP-Avenir grant, an IBISA grant and I2BC start-up funds awarded to JMS. This work was supported as part of the France 2030 programme ANR-11-IDEX-0003, which includes a research grant awarded by the interdisciplinary program OI LivingMachines@Work from University Paris-Saclay to JMS. The I2BC High-throughput sequencing facility is supported by *France Génomique* (funded by the French National Program “Investissement d’Avenir” ANR-10-INBS-09). The present work has benefited from Imagerie-Gif core facility supported by I’Agence Nationale de la Recherche (ANR-11-EQPX-0029/Morphoscope, ANR-10-INBS-04/FranceBioImaging; ANR-11-IDEX-0003-02/ Saclay Plant Sciences). W2mef parasites were obtained through BEI Resources, NIAID, NIH: *Plasmodium falciparum*, Strain W2mef, MRA-165, contributed by Alan F. Cowman. The PlasmoDB.org resource was invaluable to this work. We thank D. Noordemeer, V. Jeffers and H. Painter for critical reading of the manuscript, and K. Frenal for providing the glideosome image.

## Authors contribution

Conceptualization: JMS

Parasite culturing: LP, JMS, LM

Single cell experiment: LP, JMS,YJ

Data analysis: DN, CH, JMS

Writing: JMS

## Supplementary Information

**Figure S1.**
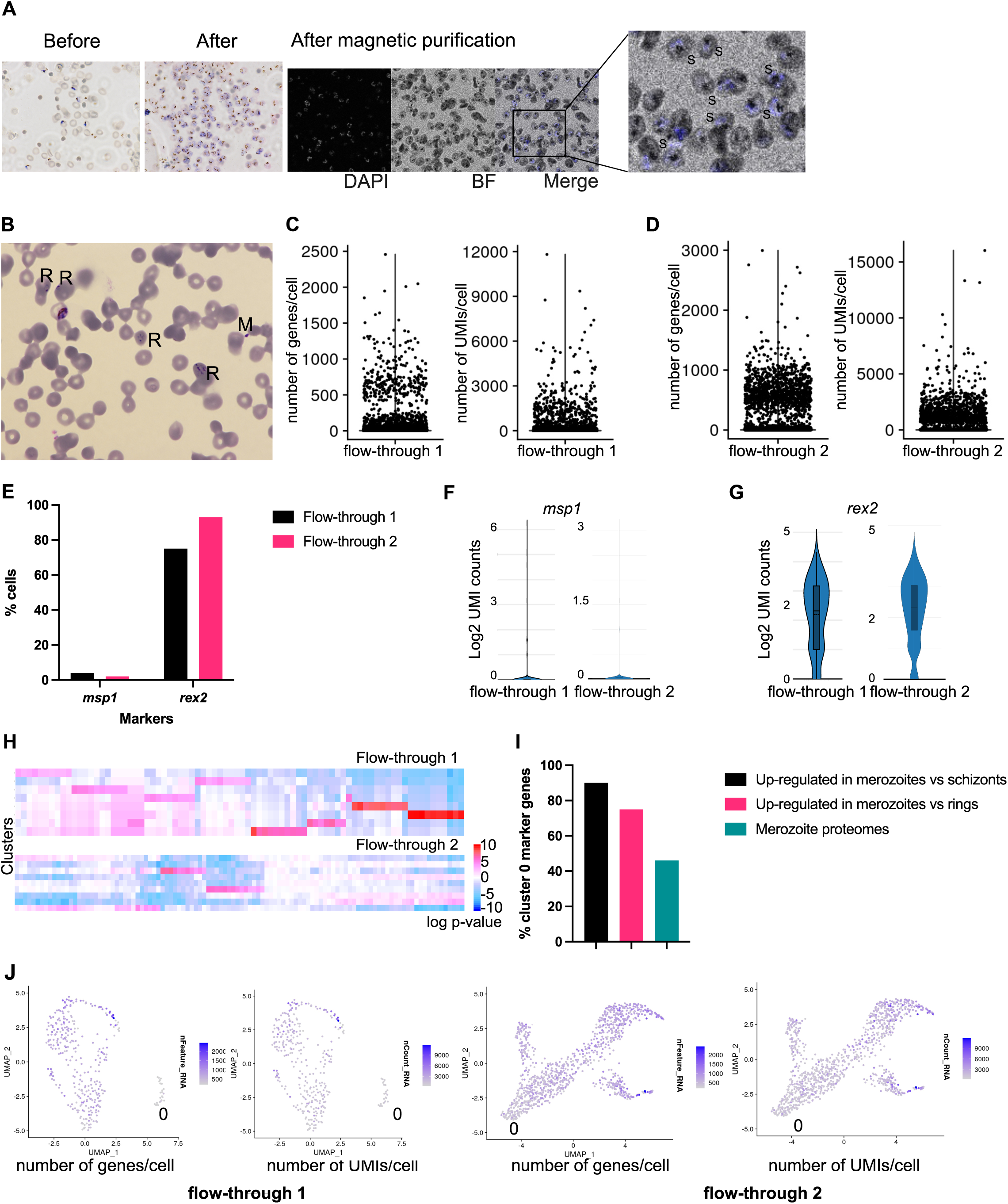
Preparation of parasites for single cell RNA-sequencing. (**A**) Giemsa-stained blood smears of schizonts before (left) and after (right) magnetic purification. DAPI-staining of the purified schizonts (S) is also shown (BF: brightfield). Higher magnification of a DAPI staining region is shown. (**B**) Giemsa-stained blood smear of the flow-through fraction of the parasite culture 1 used for single cell RNA-sequencing showing the presence of rings (R) and free merozoites (M). (**C**) Plots showing the number of genes and UMIs detected per flow-through cell of the parasite culture 1. (**D**) Plots showing the number of genes and UMIs detected per flow-through cell of the parasite culture 2. In **C** and **D**, each dot represents a cell. (**E**) Percentage of flow-through cells expressing the merozoite marker *msp1* or the ring marker *rex2*, showing that some cells express the merozoite marker. (**F**) Violin plot showing the expression level of the merozoite marker *msp1* in the flow-through cells. (**G**) Violin plot showing the expression level of the ring marker *rex2* in the flow-through cells. (**H**) Heatmap showing differential expression of 100 randomly selected genes in the different clusters of the two flow-throughs. Each row represents a cluster. (**I**) Bar graph showing how many of the marker genes of the cells of the cluster 0 are over-expressed in merozoites versus schizonts (black) or are over-expressed in merozoites versus rings (pink) according to Reers *et al*. (4), or encode proteins identified in the two merozoite proteomes (green) (13,14) (The log2 expression values determined by Reers *et al*. for each gene and which genes encode proteins detected in the proteomes is in Table S2). (**J**) UMAP representation of the flow-through cells of the two parasite cultures colored by number of genes or UMIs (reads) detected per cell, showing that cluster 0 cells express less genes and have less UMIs per cell.

**Figure S2.**
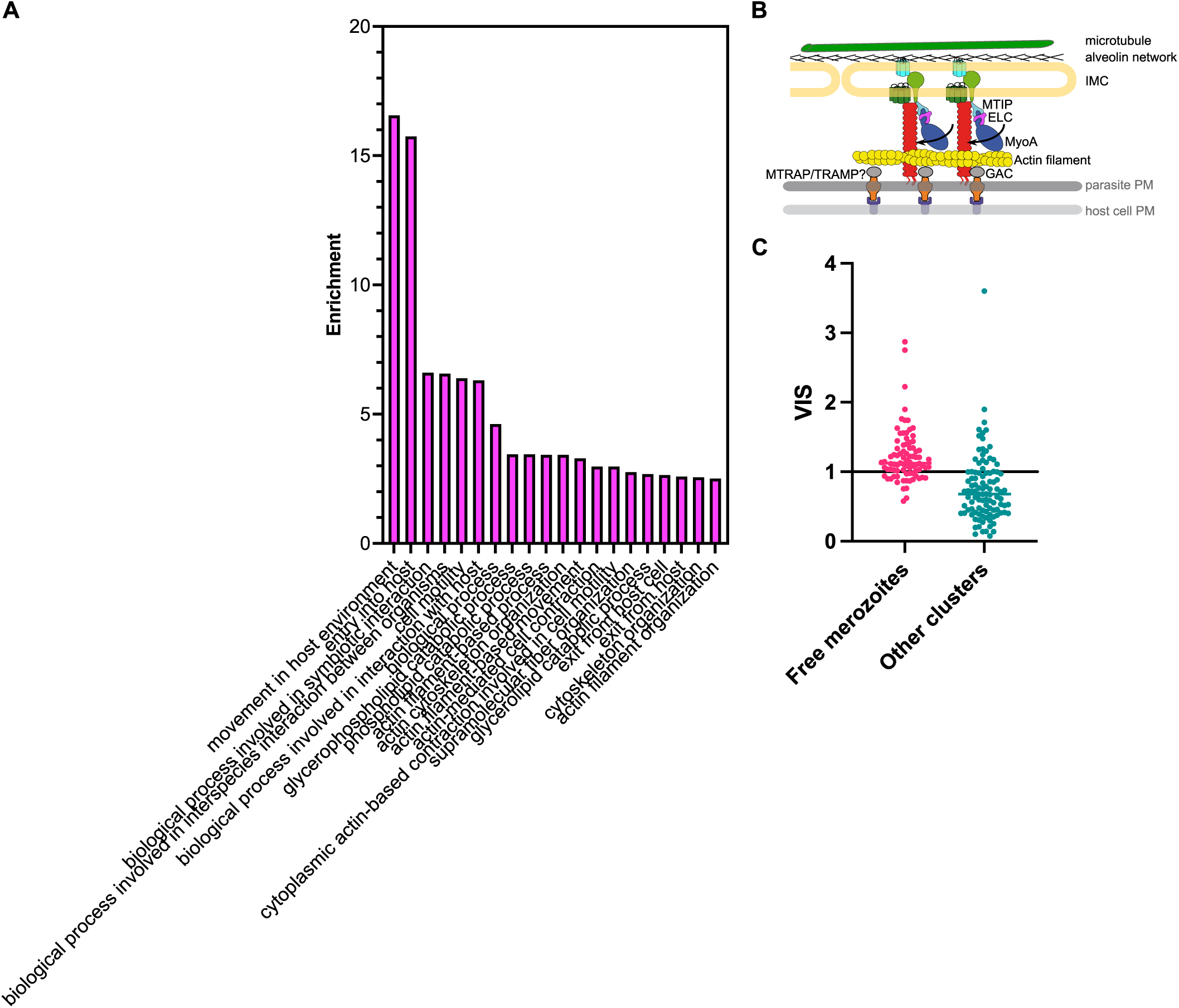
Analysis of the marker genes of free merozoites. (**A**) GO-term analysis of the marker genes of the free merozoites. (**B**) Scheme of the parasite glideosome, modified from (27), with the authorization of the authors, showing the different molecular components. The annotated proteins are encoded by genes over-expressed in free merozoites. PM: plasma membrane, IMC: inner membrane complex. (**C**) Graph showing the average variability index score (VIS) of each free merozoite gene (pink) as compared to the marker genes of other clusters (blue), as calculated by Tripathi *et al*. (8). The VIS score for each gene is in Table S2.

**Figure S3.**
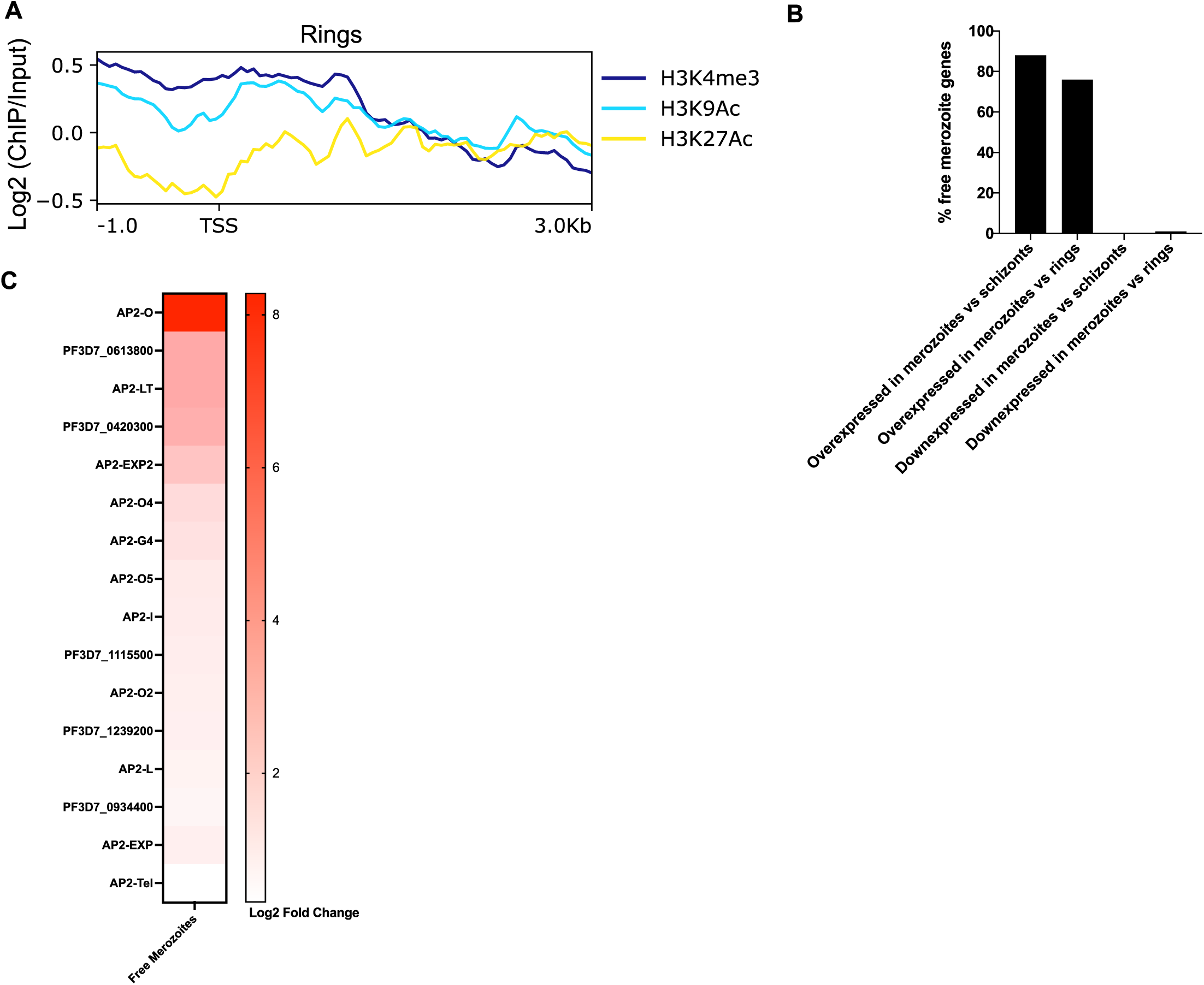
Gene expression analysis of the free merozoites. (**A**) Graph-plots showing binding of the histone marks H3K9ac (light blue), H3K27ac (yellow) and H3K4me3 (dark blue), to the promoters, transcription start sites (TSS) and gene bodies of the free merozoite genes in ring-stage parasites, as determined by Reers *et al*. (4). (**B**) Percentage of marker genes of free merozoites over-expressed in merozoites versus schizonts or rings or under-expressed in merozoites versus schizonts or rings, as determined by Reers *et al*. (4). (**C**) Heatmap showing expression of *apiap2* genes in free merozoites.

**Figure S4.**
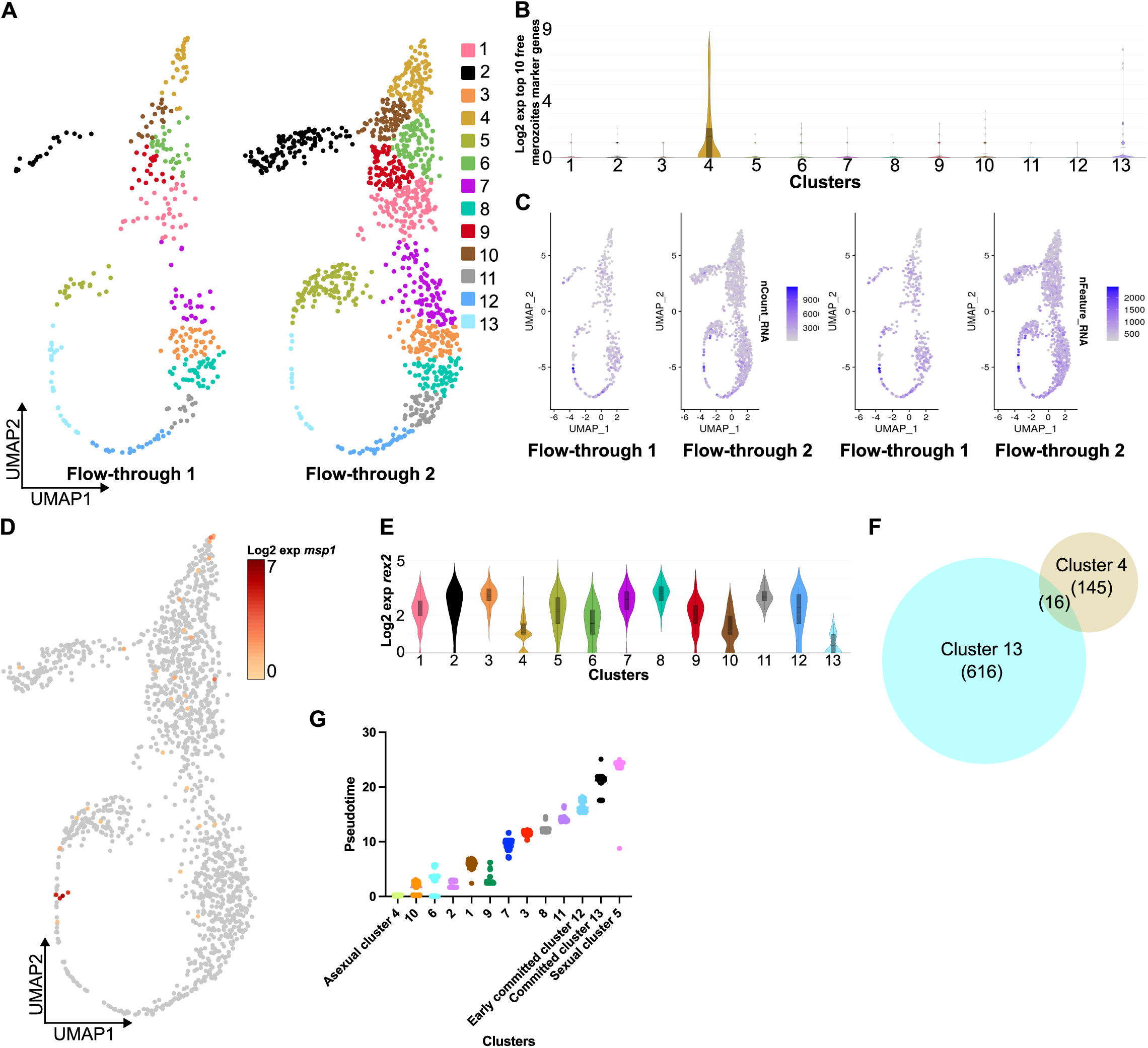
Identification of two early sexual commitment states. (**A**) UMAP representation of the Seurat clusters of the combined flow-through dataset for each flow-through sample (the number of cells in each cluster is in Table S1, the marker genes of each cluster is in Table S3, the code used is in supplementary file 3). (**B**) Graph showing higher expression of the top 10 free merozoites marker genes in clusters 4 and 13. (**C**) UMAP representation showing that some of the cells in clusters 4 and 13 contain less RNA (nCount) and encode less transcripts (nFeature). UMAP representation showing that cells in cluster 4 and cluster 13 cells express the merozoite marker *msp1* (**D**) and under-express the ring marker *rex2* (**E**), indicating that they are free merozoites. Violin plots are colored according to the cluster. (**F**) There are only 16 common marker genes between the clusters 4 and 13 and all of these are markers of free merozoites (the identity of these genes is in Table S3). (**G**) Pseudotime of the cells in each cluster as determined by Monocle. Boxes are colored according to the corresponding cluster. The pseudotime of each cell is Table S1 and the Monocle analysis is in supplementary file 3.

**Figure S5.**
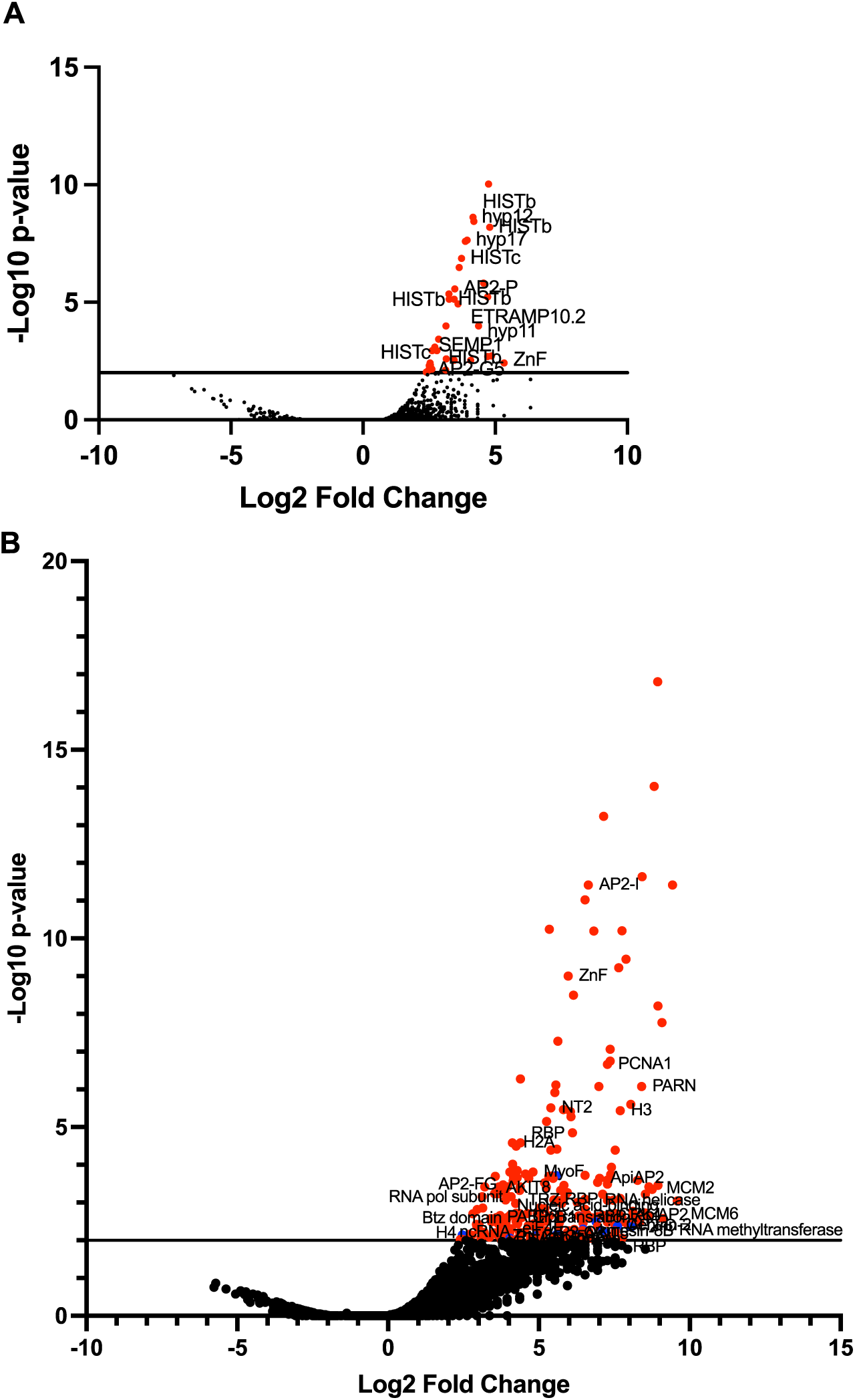
Marker genes of early committed and committed cells. Volcano plots showing in red the genes over-expressed by the early committed cells – cluster 12 (**A**), or the committed cells – cluster 13 (**B**) (the full list of genes is in Table S3). In (**B**), in blue, are free signature genes. The names of some of the genes are shown.

**Figure S6.**
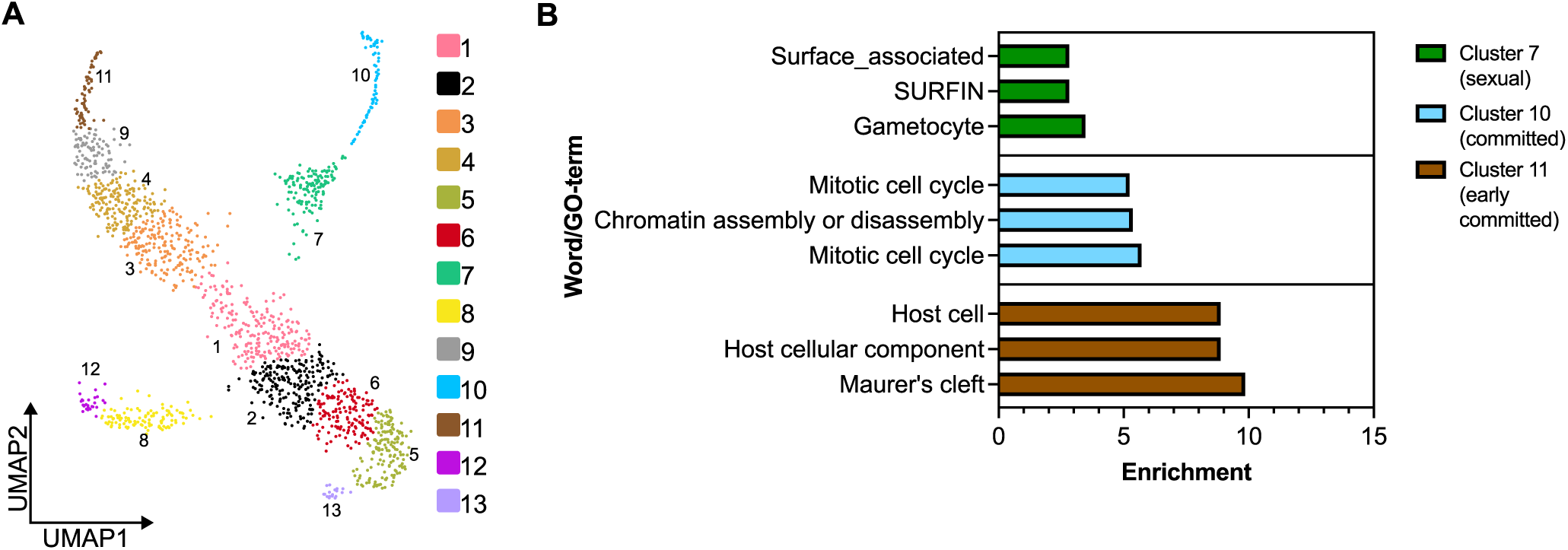
Identification of intracellular early committed and committed cells. (**A**) UMAP plot showing the Seurat clusters of the integrated dataset of the flow-throughs and intracellular cells. Cells are colored according to the cluster (the number of cells in each cluster is in Table S1 and the Seurat analysis are in supplementary file 4). (**B**) GO-term analysis of the marker genes of early committed (cluster 11), committed (cluster 10) and sexual cells (cluster 7). Bar graphs are colored according to the corresponding cluster. Only the most significant GO-terms are shown. For the complete GO-term analysis, refer to Table S4.

**Table S1.** Number of cells and reads obtained after single cell RNA-sequencing, number of cells in each Seurat cluster for the intracellular and flow-through cells.

**Table S2.** Gene expression and GO-term analysis of the genes over-expressed on the flow-through SEURAT clusters (it includes both flow-throughs). It includes the list of marker genes of free merozoites. It includes the peak transcription, the ATAC-seq peak and the VIS analysis, the comparative ChIP-Seq analysis and the comparison with the Reers *et al*. data, as well as with the merozoite proteomes. The AP2-LT ChIP-Seq data is from Bonnell *et al*. and Shang *et al*., the AP2-I ChIP-Seq data is from Santos *et al*. and Josling *et al*., the GCN5 ChIP-Seq data is from Lucky *et al*. and Tang *et al*., the transcription and stability peak time is from Painter *et al*., the ATAC-Seq data is from Toenhake *et al*., the VIS data is from Tripathi *et al*., the merozoite vs ring/schizonts log2 expression data is from Reers *et al*., the merozoite proteomes data is from Khan *et al*. and Kumar *et al*.

**Table S3.** Gene expression and GO-term analysis of the genes over-expressed on the integrated flow-through SEURAT clusters. It includes the list of marker genes of asexual, early committed, committed and sexual cells, the pseudotime analysis, the comparison with other sexual commitment scRNA-seq datasets and the comparative ChIP-Seq analysis. The AP2-LT ChIP-Seq data is from Bonnell *et al*. and Shang *et al*., the AP2-I ChIP-Seq data is from Santos *et al*. and Josling *et al*., the AP2-G ChIP-Seq data is from Josling *et al*., the MORC ChIP-Seq data is from Singh *et al*. 2024, Singh et al. 2025 and Chahine *et al*., the AP2-P ChIP-Seq data is from Singh *et al*. 2025 and Subudhi *et al*., the sexual commitment data is from Dogga *et al*. and Mohammed *et al*..

**Table S4.** Gene expression and GO-term analysis of the genes over-expressed on the integrated dataset of the intracellular cells and two flow-throughs SEURAT clusters.

**Supplementary file 1 – Code used to analyze the flow-through cells of parasite culture 1. It includes the Seurat analysis.**

**Supplementary file 2 – Code used to analyze the flow-through cells of parasite culture 2. It includes the Seurat analysis.**

**Supplementary file 3 – Code used to integrate the two flow-through datasets. It includes the Seurat and Monocle analysis.**

**Supplementary file 4 – Code used to integrate the two flow-through datasets with the intracellular cells. It includes the Seurat analysis.**

